# Two types of response inhibition failures in humans

**DOI:** 10.64898/2026.08.07.743566

**Authors:** Stein F. Acker, Jan R. Wessel

## Abstract

Response inhibition is a key control function that allows humans to perform safe, goal-directed behaviors. The dominant behavioral-computational model holds that response inhibition fails when the inhibition process is too slow to intercept unwanted movements. However, many have hypothesized that some inhibitory failures instead result from a failure to launch the inhibition process altogether. Since no method exists to identify individual ‘trigger failure’ (TF) trials, they are hitherto a largely hypothetical phenomenon. We combined Bayesian process-mixture modeling with likelihood ratio testing and jackknife resampling to identify TF trials in 253 humans. We find that TF indeed represent a separate category from other inhibitory failures. Unlike other inhibitory failures, TF do not result from premature responses. Furthermore, TF lack a stop-signal P3 event-related potential, a neural index of response inhibition. Surprisingly, TF do not result from perceptual/attentional lapses. Thus, TF are a qualitatively distinct class of executive failure during response inhibition.

## Introduction

The ability to stop and cancel unwanted or inappropriate actions is key to adaptive behavior. It allows humans and other cognitive agents to avoid adverse consequences (e.g., by stopping to walk into the street when we suddenly see a car), pursue complex internal goals (e.g., by overcoming preexisting habitual responses), and navigate everyday social environments (e.g., by inhibiting impulsive behaviors).

Measuring response inhibition in the laboratory is difficult. Conflict-based paradigms or Go-Nogo tasks are sometimes used, but are marred by the fact that response inhibition failures in these tasks can be avoided by responding more slowly ^1^. Therefore, for more than forty years, the gold-standard laboratory measurement of response inhibition has been the stop-signal task ^2–4^. In the stop-signal task, an initial Go-signal is followed, with some delay, by a second signal – the stop-signal. Varying the stop-signal delay (and, often, yoking it to ongoing behavior) countermands these slowing strategies. Moreover, it allows the calculation of a reaction time measure (stop-signal reaction time, SSRT) - a measure of the speed of the response inhibition process. This seems counterintuitive, since successful stop-signal trials do not feature an overt response. The key breakthrough in this regard was Logan and colleagues’ formulation of a behavioral-computational process model – the horse race model^3^. The horse race model infers the speed of the response inhibition process from the reaction time distributions and response inhibition failure rates at different stop-signal delays ^5^. Computationally, it is implemented as a race between two noisy accumulators – one representing the motoric ‘go’-process that races to execute the response, and one representing the inhibition process that races to cancel the response ^6^. The success of stopping is determined by which accumulator reaches a (fixed or variable ^7,8^) threshold first.

Therefore, a key corollary of the horse race model is that on individual trials, successful response inhibition depends solely on the speed of the inhibition process (triggered by the stop-signal) relative to the speed of the go-process (triggered by the go-signal). According to the horse race model, failures of response inhibition occur when the inhibition process is too slow to ‘win the horse race’ against the go process.

However, a long-standing problem for the stop-signal paradigm and the horse race model is the potential existence of stop-signal trigger failure trials ^5^. Trigger failure trials are instances in which response inhibition fails not because the stopping process was too slow, but because it was never initiated to begin with ^9,10^. These trials violate the assumptions of the classic horse race model. In turn, they render estimates of the speed of stopping via SSRT to be highly inaccurate^11^. The existence of these trigger failure trials has been long theorized (e.g., ^12^) and is perhaps most strongly indicated by non-zero intercepts of subjects’ inhibition functions – in other words, by the fact that inhibition failures sometimes occur even when the stop-signal is presented at the same time as the go-signal. Since SSRT is typically much shorter than regular reaction times, this should be theoretically impossible. However, identifying trigger failure trials empirically has been a challenging problem. Early work could not identify reliable metrics to quantify even the mean rate of trigger failures in stop-signal behavior (e.g., ^9^). Arguably the most notable progress in recent years was the formulation of a hierarchical Bayesian process model (*Bayesian estimation of ex-Gaussian stop-signal reaction time distributions*, BEESTS; ^13,14^) as a method to estimate the overall rate of trigger failures in each individual. BEESTS uses an ex-Gaussian processing model and a hierarchical mixture-likelihood approach to estimate the subject-averaged trigger failure rate from a given individual’s behavior.

Research using BEESTS to identify subject-averaged trigger failure rates has been tremendously successful in generating new potential insights into response inhibition. For example, trigger failure rates correlate more strongly with self-reported impulsivity than SSRT ^15^. Trigger failures rates are also significantly increased in ADHD ^16^, as had been long hypothesized ^12^. Similar increases in trigger failure rates have been reported in Schizophrenia ^17^. Trigger failure rates have also been correlated with electrophysiological ^18^ and BOLD measurements ^19^ of response inhibition, and are increased in subjects with brain lesions in regions previously associated with SSRT ^20^. Finally, trigger failure rates are increased during mind wandering ^21^ and decreased when successful stopping is rewarded ^22^.

However, despite the tremendous promise shown by this type of correlational work, it is important to realize that – like SSRT – trigger failures are hitherto largely a hypothetical construct. Indeed, influential work has questioned whether they exist to begin with^23^. The main challenge is that it is hitherto impossible to distinguish individual instances of trigger failures from other failed stop-trials within a given subject. Rather than correlating mean rates of trigger failures across subjects with other parameters (which is possible using existing models, see above), distinguishing individual instances of trigger failures from other response inhibition failures would allow a test of several highly specific hypotheses regarding the purported qualitative difference between trigger failures and other inhibitory failures. We will highlight three key hypotheses in the following.

First, unlike ‘regular’ response inhibition failures, which are due to the insufficient speed of the – properly triggered – inhibition process relative to the go-process, the speed of the go-process should not influence the success of stopping on trigger failure trials (as it does not matter how slow or fast the go process is if no inhibition is ever initiated). In other words, reaction times on trigger failure trials should not be faster than those on regular go-trials. This is in stark contrast to the predictions of the horse race model, according to which failed stop-trials oversample the fast portion of the go-reaction time distribution, since trials with faster RTs are more likely to lead to the stop-process ‘losing the horse race’. Indeed, finding faster failed stop-trial reaction times compared to go-trials is not just a ubiquitous finding in the literature, but a key diagnostic for the veracity of the horse race model. Thus, one clear prediction of the trigger failure model is that trigger failure trials should feature slower responses than ‘regular’ inhibitory failures, and should not be faster than go-trial reaction times.

Second, trigger failures should lack some of the brain activity associated with the response inhibition process (i.e., the activity observed on successful stop-trials). In contrast, the horse race model predicts that both successful and failed stop-trials should feature similar degrees of activity related to response inhibition, but that failed stop-trials should feature a slower unfolding of that activity. Indeed, the stop-signal P3 event-related potential ^24^ shows that exact property: it begins significantly later on failed compared to successful stop-trials, with little differences in amplitude ^25–27^. Thus, a second clear prediction is that trigger failure trials should feature a reduced or absent stop-signal P3, compared to both successful stop-trials and compared to other failed stop-trials.

Finally, many have hypothesized that trigger failures result from failed attentional detection of the stop-signal, rather than from an inhibitory control deficit ^16,18,20,28,29^. This would imply that trigger failure trials, compared to other failed stop-trials, should show differences in signatures of perception of, or attention to, the stop-signal. Candidate signatures are early perceptual brain activity after stop-signals (in the case of visual signals, the posterior visual N1 component, ^30^), the subsequent attention-related N2 ^31^, or pre-stimulus posterior α power ^32^.

Because no method exists to identify individual trigger failure trials, these hypotheses remain hitherto untested. In the current study, we achieve this by distinguishing individual instances of trigger failure trials from other inhibition failures. We did this using two approaches, which yielded highly comparable parameter estimates. First, we applied a jackknife resampling procedure to the trigger failure parameter values from the existing *Ex-Gaussian Stop-signal* (EXG-SS) model. EXG-SS is an implementation of BEESTS that is part of the *Dynamic Models of Choice* package (DMC; ^33^), and allows a quantification of the subject-averaged trigger failure rates. On each iteration, one specific stop-trial was held out. The resulting change in the subject-average trigger failure parameter was then treated as an estimate of the trial-wise parameter (see ^34^). In a second approach, we used likelihood ratio testing to derive an analytical solution that calculated the trial-to-trial trigger failure probability from the subject-level parameters estimated by EXG-SS. We applied both modeling approaches to 253 datasets of human subjects performing the stop-signal task while also undergoing whole-scalp EEG.

We then tested whether these trial-wise trigger failure probabilities were related to the trial-wise event-related EEG response to the stop-signal on failed stop-trials. Indeed, several spatiotemporally coherent clusters of stop-related EEG activity showed highly significant correlations with trigger failure probability. This shows that trial-to-trial variation in trigger failure probability is meaningfully related to specific patterns of trial-wise brain activity. We then investigated the subject-wise distributions of the estimated trial-wise trigger failure parameter values via Gaussian mixture modeling. We found that these values do not follow a single-mixture distribution, which would indicate a single generative process. Instead, these parameter values showed a bimodal distribution indicative of two separate mixture processes (one that reflects trigger failures and one that does not). This allowed us to classify each subjects’ failed stop trials into trigger failure trials and non-trigger failures to test the three hypotheses about trigger failures outlined above: whether trigger failure trials show response time distributions that are more similar to go-trials, whether they show a reduced fronto-central P3, and whether they show reduced neural indices of perceptual or attentional detection of the stop-signal.

## Methods

### Participants and task

Data were taken from published EEG studies of healthy adult undergraduates at the University of Iowa performing the stop-signal task ^35,36^, which were also included in a recent meta-analysis ^27^. These data are publicly available at https://osf.io/v3a78/ and a detailed description of the demographics can be found in the above-mentioned meta-analysis. All research was approved by the University of Iowa Institutional Review Board (IRB 201511709) and performed in accordance with the Declaration of Helsinki.

The task was a standard visual version of the stop-signal task. A detailed description is available in Wessel (2020). In short, trials began with a fixation cross (500ms duration), followed by a white leftward or rightward arrow (go-signal). Participants were instructed to respond as fast and accurately as possible to the arrow using their left or right index finger (the respective response buttons were q and p on a QWERTY keyboard). On one-third of trials, a stop-signal occurred (the arrow turned from white to red) at a delay after the go-stimulus (the stop-signal delay, SSD). The SSD, which was initially set to 200ms, was dynamically adjusted in 50ms increments to achieve a p(stop) of 0.5: after successful stops, the SSD was prolonged; after failed stops, it was shortened. This was done independently for leftward and rightward go-stimuli. Trial duration was fixed at 3000ms. Six blocks of 50 trials were performed (in total, there were 200 go and 100 stop trials).

Of the 255 subjects in the original dataset, one was excluded due to an excessive number of anticipatory responses (70 response times under 200ms), while another was excluded after failing 98 out of 100 stop trials, leaving a final sample of N=253.

### Bayesian mixture modeling

We implemented a Bayesian mixture model that is based on the existing EXG-SS model in DMC ^33^. Instead of using the original EXG-SS model in DMC, we implemented a more computationally efficient model to facilitate the computationally intensive jackknifing approach (see below). This was achieved by implementing the EXG-SS functions `log.prior.dmc()`, `h.log.likelihood.dmc()`, and `likelihood.dmc()` in C++ and by reorganizing the `likelihood.dmc()` function to eliminate redundant calculations. C++ was chosen due to its robust bindings in R through the Rcpp package^37^. Once ported into C++, calculations were performed using Armadillo ^38^ and the GNU Scientific Library ^39^. These libraries were made compatible with Rcpp through RcppArmadillo and RcppGSL, respectively.

Given parameters θ = [µ, σ, τ] and time in seconds t, the EXG-SS model represents the likelihood that a runner finishes its race at t through the ex-Gaussian probability density function f(t | θ). It represents the probability that a runner has not yet finished its race at time t through the ex-Gaussian complementary cumulative distribution function F̅(t | θ). Additionally, it accounts for probability p_gf_ of a go failure, in which case the go runner never begins, as well as probability p_tf_ of a trigger failure, in which case the stop runner never begins. Thus, it represents the likelihood of any response and the probability of no response through the following equations, respectively:

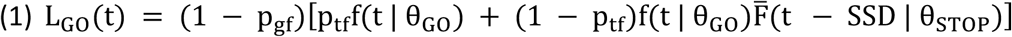

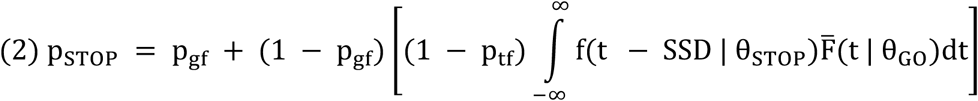

In the original EXG-SS model, the calculation of L_GO_(t) is implemented as written above. However, this function calculates f(t | θ_GO_) twice. We removed this redundancy by pre-calculating f(t | θ_GO_) and using its cached value in both terms.

The calculation of p_STOP_is more complex. Because the precise value of t cannot be determined, the likelihood of successfully stopping is integrated over all possible values of t. However, this integral is mathematically intractable, so it is necessary to perform numerical integration. In the original EXG-SS model, this is performed through Gauss-Kronrod adaptive quadrature using R’s built-in `integrate()`function for each value of SSD present in each condition. This numerical integration approach is very computationally expensive. Additionally, because EXG-SS encodes different go signals as different conditions, the `integrate()` function may be called multiple times to generate the same result for the same SSD. We streamlined this in two ways: first, we ran GSL’s more efficient Gauss-Kronrod adaptive quadrature algorithm `gsl_integration_qagi()` only once for each value of SSD found in the dataset. Second, we performed many simple calculations between constants ahead of time to avoid their repeated calculation during the integration process.

For validation purposes, we ran a hierarchical analysis on five participants with probit-transformed trigger failure and go failure parameters using both the original EXG-SS implementation, as well as our optimized model. This analysis was performed 20 times in parallel for each architecture on an Intel Core i5-13500 CPU, which launched each analysis as a single-threaded process. Elapsed time for each was measured in seconds starting at the individual parameter estimation step and ending after hierarchical analysis had produced 400 samples. Doing so allowed us to compare the outputs from each model to test whether there was a statistically significant deviation in the extracted parameters. (The resulting parameters are not expected to be identical, since the parameters in each model are extracted via Differential Evolution Markov chain Monte Carlo simulations. Thus, even running the same model twice will give slightly different results). It also allowed us to compare the computational efficacy of both implementations.

Prior to modeling, any trial with an RT below 200ms was excluded (15 trials across the whole sample) as a potential anticipatory response. Prior parameters were produced according to a truncated normal distribution with mean values set to 0.5, 0.1, 0.1, and −1.5 for mu, sigma, tau, and trigger/go failure rate values, respectively, and all SDs set to the absolute value of the mean. Upper and lower truncation values were 0 and 4.0 for all values except trigger/go failure rates, which were set to −10.0 and 0. in the hierarchical step. For hyper-priors, mu values were generated as above while sigma values were generated as a gamma distribution with shape and scale of 1.0 and 1.0, respectively. The probability of a migration step was set to 5%, and this model ran until the multivariate proportional scale-reduction factor had reached <1.1.

Performing hierarchical parameter estimation of eight parameters for five participants yielded 40 separate comparisons, with each comparison consisting of 20 replicates using both the original EXG-SS model and our new implementation. Differences between parameters were assessed using per-comparison t-tests with FDR correction for multiple comparisons. All of the 40 comparisons yielded p-values above 0.1. This shows that the two implementations produce comparable parameter estimates.

Mean (±SD) execution time of our model implementation was 1717s (±113s), while mean execution time of the original EXG-SS model was 40920s (±1732s). Thus, our implementation model demonstrated a ∼24-fold performance improvement. This enabled us to perform the computationally intensive jackknife resampling approach.

### Trial-wise trigger failure parameterization: Overview

We then implemented two potential approaches to trigger failure parameterization: one based on jackknifing, and one based on an analytical solution via likelihood ratio testing.

To assess the performance of either approach in identifying trigger failures, simulated datasets were created for all participants based on their individual parameters. Like the real data, these consisted of 200 go trials and 100 stop trials, with SSDs increasing or decreasing by 50ms based on trial outcome.

### Trial-wise trigger failure parameterization: numerical jackknifing approach

We first ran a full hierarchical analysis using data from all 253 participants three times in parallel to generate sample-level priors for the jackknifing approach. Of the three iterations, onlyone successfully converged for all participants, so the parameters generated from this analysis were used. This sample-level parameterization approach is the same as in prior studies using EXG/BEESTS and was adapted from ^20^.

Jackknifing was then performed by iteratively re-running non-hierarchical parameter estimation for each individual subject, once for each stop trial, which was removed from the dataset for that iteration. Prior parameters were generated based on a truncated normal distribution with each individual’s mean and standard deviation derived from the hierarchical analysis above. Lower and upper bounds for each parameter were 0 and 4.0, except for the trigger/go failure rates, which were again bounded at −10.0 and 0. The univariate proportional scale-reduction factor was set to 1.05 in this case to ensure greater precision. The result are probit values that express the degree of change in trigger failure probability associated with each individual failed stop-trial. To facilitate interpretation and comparison to the analytic approach (see below), these probit values were inverted, such that larger values express greater trigger failure likelihood.

### Trial-wise trigger failure parameterization: analytic likelihood ratio approach

As an alternative to the numerical brute force jackknifing method, we also used likelihood ratio testing to derive a potential analytic solution to identify trial-wise trigger failure probability from the basic parameters of the hierarchical EXG-SS model. An example of likelihood ratio testing is medical diagnostic testing. To evaluate the post-test probability p_+_(+) of a disease given positive diagnostic test, clinicians rely on the true positive likelihood L_+_(+) and false positive likelihood L_–_(+) of a disease, as well as on its pre-test probability p_+_ ^40^.

In the context of the EXG-SS model, trial-wise values for the true positive and false positive likelihoods of a trigger failure, as well as its prior probability, can be derived from the model parameters. In the EXG-SS model, the formula for a response at time t, L_GO_(t) relies on the following propositions in the context of a stop trial (both of which assume that a go failure has not occurred):

1. If a trigger failure has occurred, the likelihood L_GO_(t | θ_STOP_ ∩ tf) of a response at time t can be calculated as

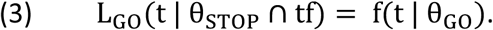
2. If a trigger failure has not occurred, the likelihood L_GO_(t | θ_STOP_ ∩ t^>^f) of a response at time t can be calculated as

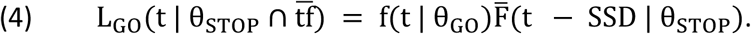

The relations between these equations are then summarized in the following formulae:

1. The positive likelihood ratio can be calculated as LR_+_(+) = L_+_(+) / L_–_(+).
2. The pre-test odds can be calculated as o_+_ = p_+_ / (1 − p_+_).
3. The post-test odds can be calculated as o_+_(+) = o_+_LR_+_(+).
4. The post-test probability can be calculated as p_+_(+) = o_+_(+) / [1 + o_+_(+)].

Given that the prior (overall) probability of a trigger failure p_tf_ is estimated during the modeling procedure, it is possible to analytically calculate the probability that any individual failed stop trial is a trigger failure using EXG-SS parameters. We can derive such a formula through the following steps:

1. The trigger failure likelihood ratio LR_tf_(t | θ_STOP_) can be calculated as

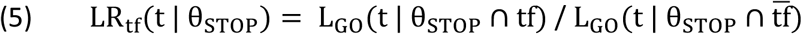 which can be simplified to

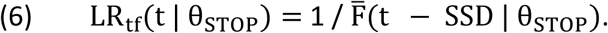
2. The prior odds of a trigger failure can be calculated from the prior probability of a trigger failure as

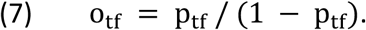
3. The odds of a trigger failure given a response at time t can be calculated as

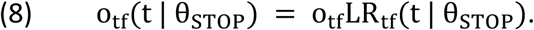
4. The probability of a trigger failure given a response at time t can be calculated as

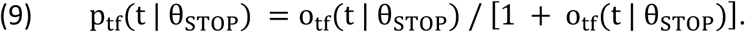

Performing all the calculations listed above yields the following formula to estimate per-trial trigger failure probability:

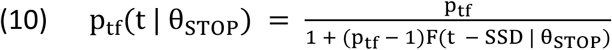

where F(t | θ) represents the ex-Gaussian cumulative distribution function, or 1 − F̅(t | θ).

### Gaussian mixture modeling of trial-wise TF parameter values

To test whether the TF values for individual subjects originate from a single generative process (null hypothesis) or from two separate processes (e.g., TF and non-TF), we performed Gaussian mixture modeling (GMM) of each subjects’ trial-wise trigger failure parameter values using a maximum number of iterations of 1,000 and a regularization factor of .0001. The number of components was set to one or two, and the goodness of fit of the resulting subject-wise models was identified using the Akaike information criterion (AIC). To ensure sufficient data for fitting, this analysis was performed only in subjects with an estimated mean TF rate of at least 10% from the full hierarchical model (N = 19). Subject-wise AIC values were then compared on the group-level using a paired-samples t-test.

### Binary classification of trigger failure parameter

Since the two-component solution provided significantly superior model fits in the Gaussian mixture model for all 19 participants, we then classified each individual trial for each subject as TF or non-TF. To this end, we sorted the failed stop-trials by the TF parameter and classified the *x*th percentile of trials with the highest trial-wise TF values as TF trials, with *x* corresponding to the estimated subject-wise TF parameter from the full hierarchical model. We then performed categorical comparisons of behavior and EEG data between the two resulting trial groupings.

We included only subjects whose data yielded at least five TF and five non-TF trials for categorical analyses. This yielded a sample of 49 subjects for both classification approaches.

We also performed all categorical analyses using a stricter criterion, according to which only trials that were classified as TF by both approaches (jackknife and analytical) were treated as TF trials. This yielded a sample of 33 subjects with sufficient trial numbers for comparison.

### Trial-wise trigger failure parameterization: Classification performance

We generated confusion matrices to assess the performance of the binary classification based on both approaches. This was done by comparing the binarization results for the simulated data with the true trial label, resulting in true positives (TF trials classified as TF trials), true negatives (non-TF trials classified as non-TF trials), false positives (non-TF trials classified as TF trials), and false negatives (TF trials classified as non-TF trials).

### EEG recording and preprocessing

EEG data were recorded and preprocessed as described in Wessel (2020) and were downloaded in preprocessed form from the existing OSF project (see above). In short, the data are from both active and passive BrainProducts recording systems with 64 channel caps, recorded at 500Hz. Preprocessing consisted of filtering (.3Hz high-pass, 30Hz low-pass), non-stereotypic artifact rejection (based on joint probability and joint kurtosis cutoffs of 5 standard deviations,^41^), and independent component analysis with subsequent removal of eye- and electrode artefacts.

### Trial-to-trial modeling of EEG activity

To test whether trial-to-trial variations in the trigger failure (TF) parameter were related to brain activity during (failed) stopping, we correlated each subjects’ trial-wise TF parameters for each failed stop-trial to the trial-to-trial EEG activity on those same trials.

Event-related EEG data at all channels were extracted from −100ms to 900ms relative to the stop-signal on all failed stop-trials. Baseline correction was performed in the 100ms pre-signal period. The resulting channel x time x trial matrix was then correlated with the trial-wise TF parameter separately for each channel and timepoint.

The resulting correlation coefficients were Fisher’s z-converted using an inverse hyperbolic tangent function to allow averaging. These values were then averaged within ten 100ms time windows spanning the entire 1,000ms epoch. The resulting subject x channel x time window matrix was then tested for significance on the group level against zero (i.e., no correlation) via paired-samples t-tests. This required 640 individual tests (64 channels × 10 time windows). Consequently, the critical p-value for significance was corrected using the Bonferroni method to .05 / 640, resulting in a critical p value of .000078125.

### Behavioral analysis

We compared median reaction time (RT) and median stop-signal delay (SSD) between TF and non-TF using paired-samples t-tests. The median RT values for both groups of trials were also compared to the median go-trial RT. We expected that non-TF stop-trials would show faster RT compared to go-trials, which is the classic pattern based on the horse race model. Crucially, we also hypothesized that TF trials would be more representative of the regular go RT distribution. Thus, we hypothesized that TF trial RT would be both a) significantly slower than non-TF trial RT and b) not significantly slower than go-trial RT.

### Event-related EEG analyses

We tested two hypotheses regarding EEG activity on TF and non-TF trials. First, we tested whether fronto-central activity, especially the P3, would be reduced on TF compared to non-TF trials. Second, we tested whether TF and non-TF trials differed with regard to a series of established markers of visual-perceptual and attentional processing.

Three ERPs were investigated, the fronto-central P3 (at electrode FCz), the fronto-cental N2 (at electrode FCz), and the posterior visual N1 (at electrode Oz). Event-related EEG data were extracted from −100ms to 900ms relative to the stop-signal and averaged separately for TF and non-TF trials. Baseline correction was performed in the 100ms pre-signal period. The resulting ERPs were tested for significant differences using sample-to-sample t-tests. The resulting vector of 500 p-values (one per sample point in the 1,000ms epoch sampled at 500Hz) was corrected for multiple comparisons using the false-discovery rate (FDR ^42^) method to achieve an overall critical p-value of .01.

Furthermore, we tested whether pre-stimulus α power differed between TF and non-TF trials. Time-domain EEG data was converted to time-frequency power in the α band via a Hilbert transform of the bandpass filtered data between 8 and 12 Hz. From the resulting complex time series, signal power was extracted by squaring the absolute value. This power time-series was then averaged in the 100ms time period leading up to the stop-signal and averaged separately for TF, non-TF, and successful stop trials. We chose a relatively short time window prior to the stop-signal due to the presence of the preceding stop-signal. Condition differences were tested using paired-samples t-tests.

## Results

### Jackknifing and likelihood ratio testing produce highly similar parameter estimates

While the two parameterization procedures are fundamentally different, the resulting parameter estimates were highly correlated between the two approaches (mean r = .51, p < 9.5 * 10^−47^). This agreement was even higher when the analysis was restricted to subjects with trigger failure rates of at least 5% (mean r = .85, p < 6.7 * 10^−25^). This shows that the limited disagreement that existed between the two approaches was due to the lower parameter correlations in subjects with very few trigger failure trials. (In those subjects, most trial-wise trigger failure parameter estimates will reflect noise around zero. Thus, correlations are expected to be low.) Together, the high convergence of parameter estimates from both methods suggests that they both capture the same aspect of behavior.

### Trigger failure probability is highly correlated with trial-to-trial brain activity

Testing the association between trial-wise EEG activity on failed stop-trials and the trial-wise trigger failure parameters across all 253 subjects revealed highly significant associations (critical p < .000078; ***Figure 1***). For both approaches (jackknife and likelihood-ratio) TF parameter values were significantly correlated with fronto-central EEG activity in the 300-400ms window after the stop-signal. In both cases, higher trigger failure likelihoods were associated less positive EEG activity at fronto-central electrodes. The likelihood ratio approach furthermore identified several weaker clusters of correlation, which were not significant in the jackknife-based analysis (see ***Figure 1***, bottom).

**Figure 1.**
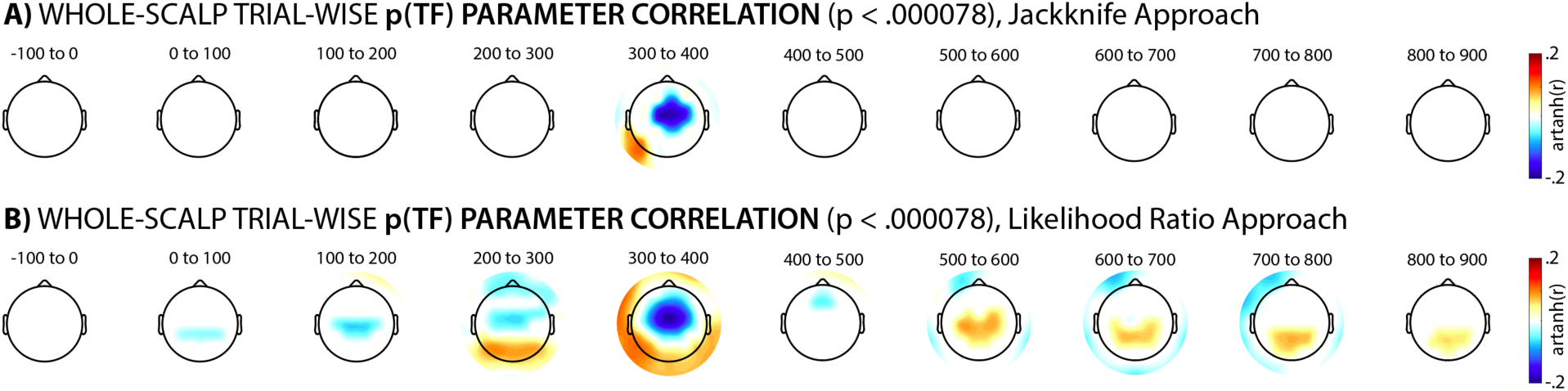
Trial-wise correlations between single-trial EEG activity at each channel and the TF parameter derived from the jackknife approach (A) and the likelihood ratio approach (B), respectively. Correlations are expressed in mean Fisher’s z-transformed correlation coefficients for each window. The resulting topographies are thresholded for significance at a critical p-value of p < .05 / 640, which accounts for the number of comparisons (64 channels times 10 time windows).

This shows that the TF parameters extracted using both approaches are related to spatiotemporally coherent clusters of EEG activity on the same trial, with converging evidence showing that trials with higher trigger failure values feature less fronto-central EEG activity.

### Gaussian mixture modeling shows that failed stop-trials are produced by two separate processes

We then tested whether the trial-wise TF parameter values show a bimodal distribution indicative of two generative processes (TF and non-TF). The trial-wise distributions of the TF parameter for the 19 subjects with sufficient trial numbers for gaussian mixture-modeling are depicted in ***Figure 2***. The AIC indicates superior fits for the two-component solution in all 19 cases for the likelihood ratio approach, and a significant group difference in AIC between the one- and two-component solutions (t(18) = 8.1962, p < 10^−6^, d = 2.5027). ***Figure 2*** depicts the best fitting per-subject models for the likelihood ratio approach, which expresses its parameters in readily interpretable probabilities (compared to the probit output from the jackknife approach). The probit parameter values derived from the jackknifing approach also favored a two-component mixture model (t(18) = 3.5897, p < 0.005, d = 0.33684).

**Figure 2.**
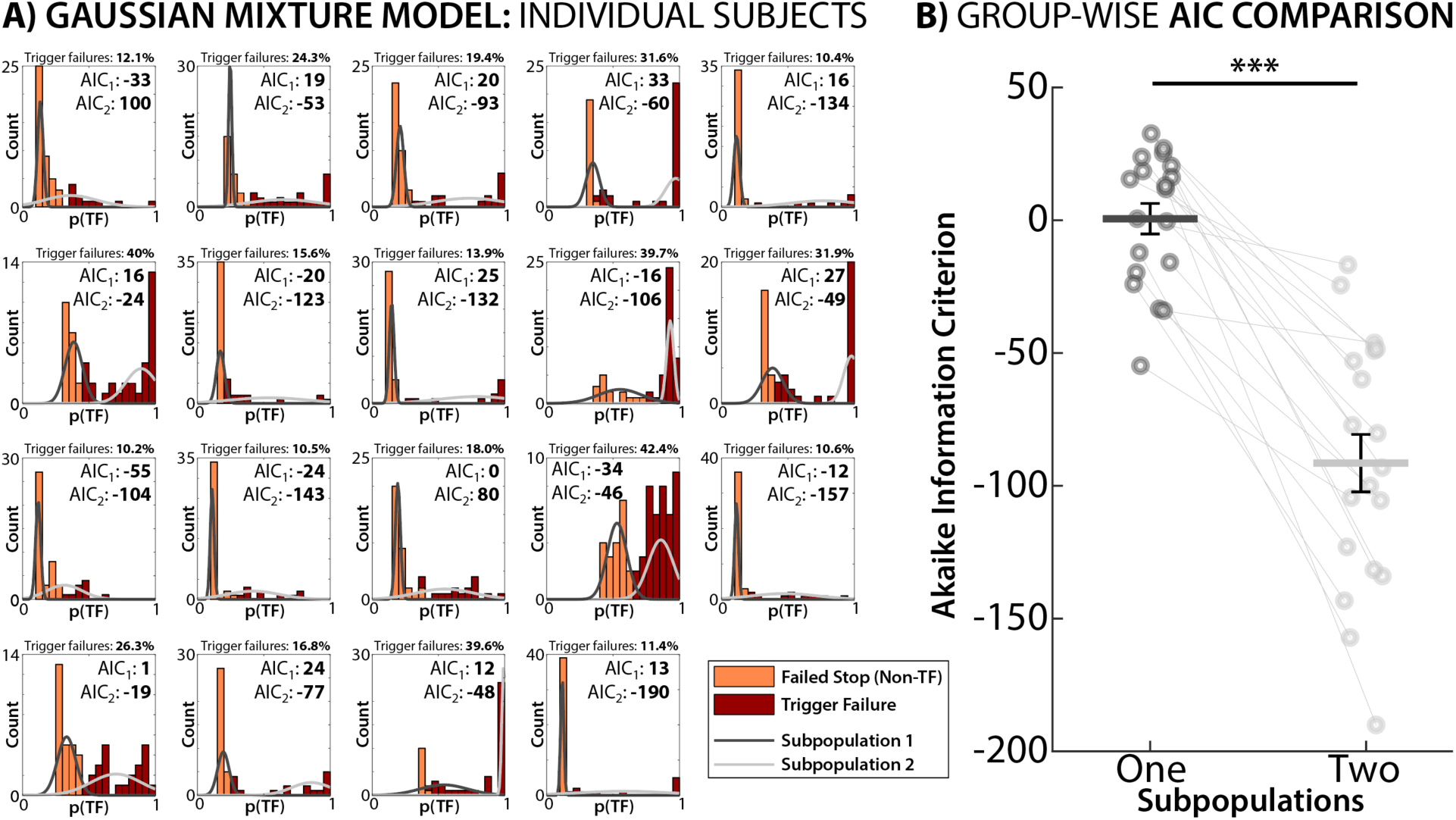
Results of the gaussian mixture modeling for the trigger failure probability distributions derived from the likelihood ratio approach in all subjects with at least 10% trigger failures. A) Parameter histograms for each subject alongside the preferred solution (according to AIC). Yellow trials are trials that were classified as non-TF failed stop trials according to the subsequent binary classification. Red trials are trials that were classified as trigger failures. Gray gaussian curves reflect the parameters returned by the winning gaussian mixture model. p(TF) = probability of trigger failure. B) Comparison of subject-wise AIC comparisons for the one- and two-component mixture models. *** = p < .0001. Error bars reflect the standard error of the mean.

In line with the trigger failure model, this suggests that failed stop-trials are generated by two processes rather than a single one. This motivates a binary classification of trials into TF and non-TF trials (see Methods) upon which the subsequent analyses are based.

### Both jackknife and likelihood methods produce accurate and sensitive binary classifications

Compared with the true values of the simulated data, the classification accuracy of the likelihood-ratio approach was 92.53% and the accuracy of the jackknife approach was 91.89%. Both approaches were highly specific, with 95.73% specificity for the analytic approach and 95.58% for the jackknifing approach. Thus, both classifications resulted in very few false positives.

In contrast, sensitivity was lower (54.41% and 48.11%, respectively). This was because both approaches had difficulty positively identifying trigger failures on trials on which stopping would have failed either way - i.e., even if there had not been a trigger failure (see Discussion). Therefore, the results of the binary classifications below should be interpreted as comparing a) trials on which stopping failed specifically because of a trigger failure (we will simply call these “trigger failures” in the following) to b) all other failed stop trials (we will call these “non-TF trials”).

### Behavior on TF trials conforms to theoretical predictions from the TF model

Using both binary classification approaches, we compared reaction times and stop-signal delays between TF and non-TF trials in 49 subjects with at least 5 trials in both categories (***Figure 3***).

**Figure 3.**
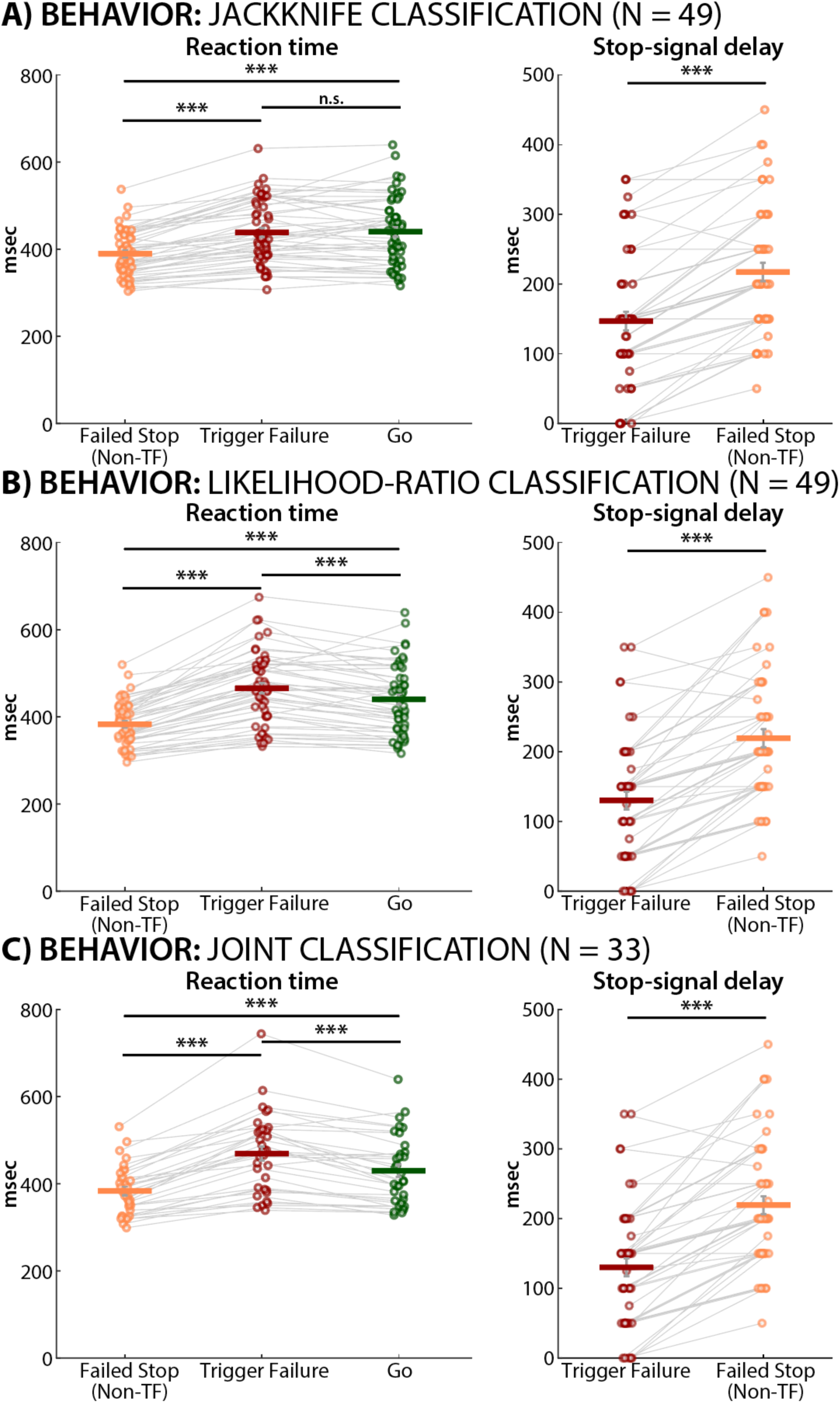
Behavioral comparisons between TF and non-TF trials based on the binary classification performed using jackknifed parameter values (A), likelihood-ratio values (B), and joint classification (C). Left panel reflects median conditional reaction times, right panel depicts median SSD. Across all three trial selections, non-TF trials showed faster RTs than go trials and TF trials. Moreover, TF trials were either statistically indistinguishable from (A) or slower than (B,C) go trials. Error bars reflect the standard error of the mean. *** = p < .0001. n.s. = not significant.

For the jackknife-based classification, regular, non-TF failed stop-trials showed the expected faster RTs compared to go trials that is expected under the horse race model (t(48) = 9.7798, p < 10^−12^, d = 0.7567). Importantly, in contrast, mean TF trial RT did not differ from go-trial RT (t(48) = 0.32442, p = 0.74703, d = 0.021766), and was significantly slower than non-TF trial RT (t(48) = 10.4337, p < 10^−13^, d = 0.77668).

For the likelihood-ratio-based classification, regular, non-TF failed stop-trials once again showed the expected faster RTs compared to go trials (t(48) = 10.5846, p = 10^−13^, d = 0.8643). Once again, TF-trial RT was significantly slower than non-TF trial RT (t(48) = 13.172, p < 10^−16^, d = 1.2202) For this classification, mean TF trial RT was actually slower than go-trial RT as well (t(48) = 4.6683, p < .0001, d = 0.3179).

The joint classification matched the pattern from the likelihood-ratio-based classification. Once again, regular, non-TF failed stop-trials showed the expected faster RTs compared to go trials (t(32) = 8.9501, p < 10^−9^, d = 0.67573). Furthermore, mean TF trial RT was slower than both go-trial RT (t(32) = 5.7217, p < 10^−5^, d = 0.46673), and non-TF trial RT (t(32) = 10.1919, p < 10^−10^, d = 1.1454).

Thus, regardless of classification approach, the behavioral results confirm core predictions of the TF model. Since TF trials result from a lack of initiation of the stopping process, they do not result from systematically faster, premature responses (as opposed to non-TF trials, which over-represent the fast part of the go-trial RT distribution, where the go-process is more likely to ‘win the horse race’). In fact, the likelihood-ratio and joint classifications suggest that TF trial RT is significantly slower than go-trial RT, though the effect sizes for these comparisons were lower (see Discussion).

We also found that trigger failures occurred more often at shorter stop-signal delays for all three classification approaches (jackknife: t(48) = 10.2563, p < 10^−12^, d = 0.76229; likelihood-ratio: t(48) = 10.2778, p < 10^−12^, d = 1.0046; joint: t(32) = 10.8969, p < 10^−11^, d = 1.2696). Once again, this is sensible, since non-TF trials should feature fewer trials at shorter SSDs (where the stop-process is more likely to ‘win the horse race’), whereas the SSD should not matter on TF trials.

### Trigger failure trials show a substantially reduced fronto-central P3

We then compared ERPs between TF and non-TF trials in subjects who had sufficient trial numbers for a trial-average ERP (N=43 for the jackknifing-based classification, N=41 for the likelihood-ratio-based classification, and N=22 for the joint classification; these subjects had at least 5 trials for both the TF and the non-TF conditions after rejection of trials with EEG artifacts). Regardless of classification approach, this analysis shows a significantly reduced (and virtually absent) fronto-central P3 waveform on TF trials compared to non-TF trials (red vs. orange lines in ***Figure 4, left column***; p < .01, FDR-corrected). The time periods of significantly reduced stop-signal P3 were highly similar across all classification schemes (322-454ms for jackknifing, 304-458ms for likelihood ratio, 322-468ms for joint classification).

**Figure 4.**
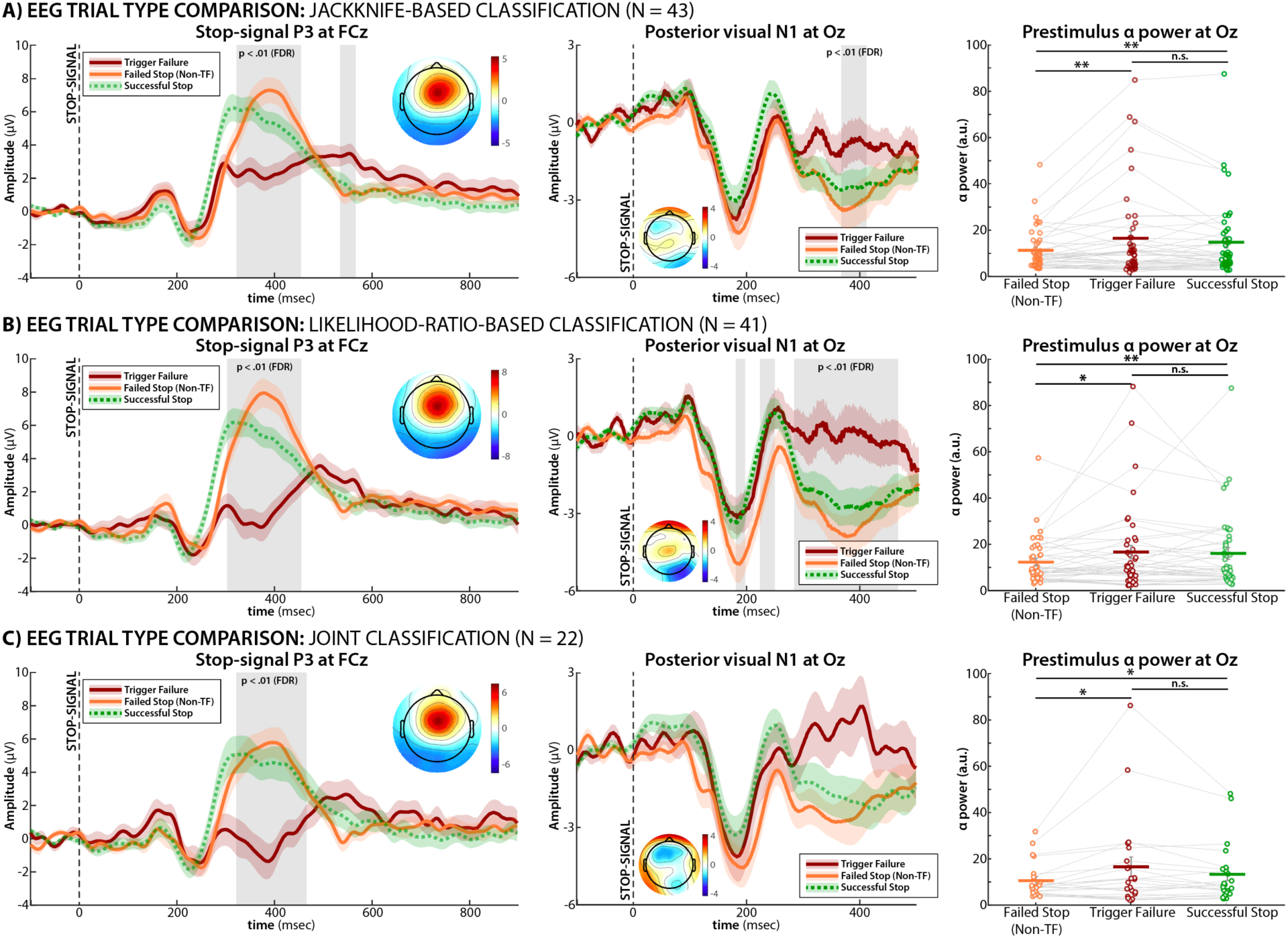
EEG comparisons between TF and non-TF trials based on the binary classification performed using jackknifed parameter values (A), likelihood-ratio values (B), and joint classification (C). The left figures shows the stop-signal locked event-related potential at fronto-central electrode FCz. The ERP time courses show that there is a significant reduction of the stop-signal P3 on trigger failure trials (gray shaded areas depicts time periods of p < .01, FDR-corrected for multiple comparisons). Voltage topographies depict the Non-TF minus TF condition difference at the peak latency of the difference wave. Shaded areas around the ERP curves reflect the SEM. The middle figures shows the stop-signal locked event-related potential at posterior electrode Oz. The ERP time courses show that there is no difference between the TF condition and successful stop trials (gray shaded areas depicts time periods of p < .01, FDR-corrected for multiple comparisons). Voltage topographies depict the Non-TF minus TF condition difference at the peak latency of the N1 (190ms). Shaded areas around the ERP curves reflect the SEM. The figures in the right column show prestimulus α band power in the 100ms leading up to the stop signal. Prestimulus α was decreased on non-TF compared to both TF and successful stop-trials across all classification approaches, with no significant differences between TF and successful stop trials. ** = p < .01, * = p < .05, n.s. = not significant. Error bars reflect the standard error of the mean.

As such, the P3 ERP shows the exact properties that would be expected from a neural index of response inhibition under the trigger failure hypothesis: Regular (non-TF) failed stop-trials and successful stop-trials differed mainly in timing, with successful trials showing an earlier P3 onset (the standard pattern in ERP studies of the stop-signal task ^25,26^). TF trials, in turn, feature a substantial reduction of the fronto-central P3.

### Trigger failures are not failures of stop-signal detection

We then tested whether there were any significant differences in known EEG markers of perceptual or attentional processing. As can be seen from the fronto-central plots (left columns in ***Figure 4***), the fronto-central N2 did not differ between the trial types in any classification scheme. The posterior visual N1 did show significant differences between TF and non-TF trials in the likelihood-ratio-based classification, but not the other two (***Figure 4, middle column***). However, this difference was driven by an increased N1 on non-TF trials compared to both TF and successful stop-trials. TF trials and successful stop-trials did not show any significant differences at any time point, in any of the classification schemes. (Note that there are later time periods of significant TF vs. non-TF difference in the jackknifing and likelihood ratio classification schemes from around 300ms onwards. These reflect dipole projections from the stop-signal P3, which dominates the voltage topography in that time range; see topographies in the left column). Finally, we compared prestimulus α power as an index of attentional state at the time of stop-signal presentation (***Figure 4, right column***). Increases in prestimulus α purportedly index different anticipatory attentional processes (e.g., target enhancement or distractor suppression, ^43,44^). Across all three classification schemes, prestimulus α activity was actually decreased on non-TF trials compared to both TF trials, rather than vice versa (jackknife: t(42) = 2.7737, p = 0.008, d = 0.34705; likelihood ratio: t(40) = 2.0358, p = 0.048, d = 0.29831; joint: t(21) = 2.1082, p = 0.047, d = 0.40205). Non-TF trials also showed decreased prestimulus α compared to successful stop-trials (jackknife: t(42) = 2.9299, p = 0.005, d = 0.27044; t(40) = 3.3907, p = 0.0016, d = 0.28249; joint: t(21) = 2.349, p = 0.0287, d = 0.26804). TF trials, on the other hand, did not differ from successful stop-trials in any classification scheme (all p > .1).

Together, these findings consistently show that there are no differences between TF and successful stop-trials with regards to established EEG markers of visual perception or attention. Differences between TF and non-TF trials were consistently attributable to differences between non-TF and successful stop-trials. This suggests that TF trials are not the result of perceptual-attentional effects (if anything, there is some evidence to suggest that non-TF failed stop-trials show altered attentional processing; see Discussion and ^45^).

## Discussion

The current study shows that humans fail to stop their actions for two qualitatively distinct reasons. In classic models of response inhibition, failures of response inhibition result solely from insufficient speed of the inhibition process relative to the unwanted movement. Here, we show that there is a categorically different type of response inhibition failure: the failure to ever initiate response inhibition. The existence of trigger failures has been long hypothesized in research on response inhibition, but had hitherto not been empirically demonstrated. The qualitative distinction between TF and non-TF trials is reflected in highly salient differences in both behavior and brain activity. Behaviorally, TF trials do not result from significantly faster responses than typical go-trials. This is in stark contrast with non-TF failures of response inhibition, which predominantly result from fast, premature responses ^3^. Neurally, trigger failures are associated with specific patterns of brain activity. Most strikingly, the stop-signal P3 event-related potential, a purported marker of response inhibition ^24,27^ is virtually absent on TF trials. Lastly, and perhaps surprisingly, TF trials did not show any differences from successful stop-trials in typical neural indicators of perception or attention. This suggests that trigger failures may not be related to attentional or perceptual factors, as often hypothesized ^16,18,20,28,29^. Instead, they appear to be specific failures in triggering the executive control process of response inhibition.

The existence of trigger failure trials as a separate type of inhibitory failure has wide-ranging consequences across the field of response inhibition. First and foremost, it poses fundamental issues for all standard methods of calculating stop-signal reaction time based on the horse race model (see, e.g., ^11^). The presence of trigger failure trials fundamentally biases the calculation of SSRT. Indeed, when TF trials are removed – i.e., when failed stop-trials only contain instances of successfully triggered, but insufficiently fast response inhibition – the actual probability of successful stopping at a specific SSRT would be higher than observed. Thus, when TF trials are not accounted for, SSRT values are systematically inflated ^11^. Of course, the severity of these calculation errors depends strongly on the overall rate of trigger failures in each dataset. In the current study, the mean trigger failure rate was relatively low (3.7%). This is typical for simple stop-signal tasks with very easily interpreted go- and stop-stimuli (see e.g., ^21^ for similar rates). However, stop-signal tasks with more complex signals (including those with simple stop-signals but more complicated go-signals) have found mean trigger failure rates as high as 22% ^18^. Even in the very simple task employed here, some participants showed trigger failure rates as high as 42%. SSRT estimates in samples with TF rates of this magnitude are likely highly inaccurate (for example, a mean TF rate of 22% led to a ∼49% increase in sample-wise SSRT in ^18^). Therefore, it is paramount that take trigger failures are taking into account in all research on response inhibition, even during basic operations such as calculating SSRT. The current work offers a method to identify specific individual instances of trigger failures. Thus, in addition to allowing more precise quantifications of SSRT (which was already possible using methods that modeled subject-level trigger failure rates, such as BEESTS), the current work will also allow future investigators to study these two trial types in separation. This will enable fundamentally new insights into the neural and physiological underpinnings of response inhibition (see below).

In addition to these fundamental implications for basic response inhibition research, the existence of TF as a categorically distinct response inhibition failure has wide-ranging consequences for the interpretation of existing and future work on the topic. For example, while some early studies have found an association between SSRT and self-report measures of impulsivity (^46^; see also: ^47^), more recent work has shown that these associations can be weak ^48^, thus questioning the utility of the stop-signal task to study real-world behaviors. The current work suggests that these studies of individual differences needs to account for different types of response inhibition failures (e.g., ^15^), which may differentially relate to real-world scenarios. The current work demonstrates a method to study these types of response inhibition failures in separation, which could substantially improve the predictive value of laboratory models of response inhibition for real-world impulse control. One can easily conceive real-world impulse control behaviors that may more strongly relate to the ability to trigger response inhibition (e.g., refusing a drink at a bar), while others may be more strongly related to its speed (e.g., checking a swing in a baseball game). The same is true for clinical investigations as well. For example, some have proposed that response inhibition is the “primary deficit” observed in attention-deficit-hyperactivity disorder ^49 50 51^. Indeed, people with ADHD consistently show elongated SSRT. However, a recent study that quantified subject-level TF rates found that this elongation might primarily be attributable to higher TF rates in the ADHD group ^16^, with similar findings reported for schizophrenia ^28^. The approach in the current work enables investigations of the mechanisms underlying response inhibition in these populations for TF and non-TF trials separately, and thus will likely contribute to novel insights into the underlying mechanisms.

In the current work, we found clear differences in EEG signatures between TF and non-TF trials. Both the trial-to-trial correlation analysis across the whole sample and the categorization approach in the subsample of subjects with sufficient TF rates provided converging evidence that trigger failure trials feature substantial reductions in fronto-central stop-signal P3 activity. This ERP is the main purported EEG index of the response inhibition process ^24,25,27^. As such, it is sensible that failures to trigger response inhibition altogether would lack this signature. In contrast, TF trials did not show any reduced EEG activity related to perceptual or attentional processing (posterior visual N1, fronto-central N2, pre-stimulus α power) compared to successful stop-trials. When TF and non-TF trials differed, it was due to differences between non-TF failed stop-trials and successful stop-trials. Most notably, non-TF trials consistently featured reduced pre-stimulus α-band activity compared to both TF trials and successful stop-trials (whereas TF and successful stop-trials did not differ). This is somewhat unexpected, given that many (including us, ^16,18,20,28,29^) have tentatively speculated that trigger failures may result from an impaired detection of the stop-signal. The results of the current work instead point in another direction: that trigger failures occur *despite* the successful detection and perception of the stop-signal. This suggests that they represent a distinct failure to initiate executive function. It also suggests that the reduced speed of response inhibition that leads to inhibitory failure on non-TF trials could be partially explain by differences in attentional processes prior to the stop-signal. However, definitive tests of these hypotheses require future study combining the current approach with custom-designed experimental paradigms optimized to study attentional processes.

As such, the current approach to TF quantification will be highly useful for future work aiming to elucidate the (neuro)physiological mechanisms that distinguish TF and non-TF trials. Response inhibition research using the stop-signal task features a long history of investigations using fMRI ^52–54^, TMS ^55,56^, motor physiology ^57,58^, and other methods ^59,60^. The parameterized single-trial TF values extracted using the current approach can be readily used to model the trial-to-trial BOLD response and test whether different brain areas and circuits are involved in either type of response inhibition failure. The binarization approach would also very easily enable contrasts of BOLD activity within each subject. This would be a logical next step from already existing recent studies that have used the subject-level mean TF rates in population-wide correlations ^19^, and those that have found elevated TF rates in certain types of brain lesion patients ^20^.

Finally, we here used two fundamentally different approaches towards quantifying the trial-wise trigger failure probability, which nevertheless produced highly correlated parameter estimates. Both approaches resulted in highly accurate binary classifications and achieved especially high specificity. However, for both approaches, the sensitivity of the binary classification was much lower. This was chiefly because trigger failure trials in which response inhibition would have failed even without a trigger failure (i.e., when even a properly triggered stop runner would have ‘lost the race’) were hard to distinguish from failed stop-trials without trigger failures. This has two consequences for the categorical comparisons in the current work. First, it suggests that the true difference in EEG activity between TF and non-TF trials is even bigger than suggested by the current study. That is because the TF trials contained very little false positives, but the non-TF trials still contained some portion of trials with trigger failures. Second, this also explains why in some classification analyses, TF reaction times were not only just as slow as go-trial RTs, but even slightly slower (since TF trials that remained in the non-TF group theoretically would have faster RTs than other TF trials). Lastly, because non-TF trials still contain some portion of trials with trigger failures, our algorithms do not provide perfectly ‘clean’ samples of non-TF trials. While the TF trials selected using the current approach provide a very accurate selection of “trials in which stopping failed because of a trigger failure” (TF trials), the non-TF trials identified by these algorithms are better understood as “trials in which stopping failed because the inhibition process was – *or would have been* – too slow to intercept the response”.

In sum, we show that humans fail to stop actions for two different reasons: either because the response inhibition process was insufficiently fast (the classic assumption of the horse race model) or because they did not trigger their response inhibition process at all (the trigger failure model). We found that trial-wise TF parameter values related brain activity in meaningful and highly reliable ways and that they are generated by more than one underlying process. By classifying each subjects’ failed stop trials into TF and non-TF trials, we found converging evidence for the long-proposed TF model in both behavior and brain activity. This work provides clear-cut empirical evidence for the existence of a qualitatively different category of response inhibition failure – trigger failures – which represent outright failures to engage executive control.

